# CircRNA-disease inference using deep ensemble model based on triple association

**DOI:** 10.1101/2023.03.07.531622

**Authors:** Laiyi Fu, Hongkai Du, Ying Wang, Qinke Peng

## Abstract

Accumulating evidence indicates more and more circular RNAs (i.e. circRNAs) have played a vital role in regulating gene expression and are related to diseases through different biological procedures. Predicting circRNA-disease associations helps to conjecture possible disease related circRNA and facilitate human disease diagnosis and downstream treatment. Nevertheless, little effort was made to uncover the interaction between various diseases and circRNAs. In our work, human circRNA-disease association network is first generated using known miRNA-circRNA interactions and disease related miRNA (microRNA) information. Then we further integrated this information to compute similarity scores between human diseases and circRNAs. Here, we proposed one deep ensemble model called DeepInteract, which first used two stacked auto-encoders to explore hidden features utilizing similarity information, and adopted a 3-layer neuron network to predict the final association. Our method is capable of capturing more complex non-linear features comparing to other approaches. Our results indicate the proposed method is superior to other previous competitors. Many prediction results have been validated by some biological experiments using our model.

## I. Introduction

Following non-coding RNAs including microRNAs (miRNAs), various type of long non-coding RNAs (lncRNAs), as well as circular RNAs (circRNAs), they all represent a class of small endogenous non-coding RNAs that are highly expressed in the transcriptome of eukaryotic[1] and crucial in regulating gene expression[2]. It’s strong and convincing evidence that circular RNA dysregulation will lead to human diseases even cancers. A great amount of circular RNA disease associations are validated through biological experiments[1, 3, 4]. Specifically, miRNA is also reported to be associated widely with circRNA and gene expression, which may have a significant influence on cancer-related diseases. Using experimental results supporting human circular RNAs and diseases associations along with miRNA and mRNA, Ghosal et al. [5] constructed miRNA, circular RNA, mRNA, disease associations based on miRNA as an index, namely Circ2Traits DataBase[5]. Li et al. [6] and Jiang et al. [7] built human miRNA-disease associated database called HMDD and another database called miR2Disease. They utilized confirmed experimentally supported miRNA-disease evidence. Using these datasets and connections, we can infer verified associations between different circular RNAs and diseases by implementing ternary relations among circular RNAs, miRNAs, and diseases.

As more and more attention focused on circRNA, the recent researches have gradually uncovered the disease mechanism related with circRNA[3]. Predicting circRNA-disease association using computational models could help researchers to better infer potential relation. Current machine learning methods for predicting association including network-based methods, and other computational methods.

Those approaches that apply machine learning techniques have been widely adopted to address these issues in practice[8] and have proved to be helpful in increasing model accuracy and classification performance[9]. Those models dealing with miRNA-disease association inferring, such as support vector machine model[10], lasso regression model[11], etc. However, these machine learning based approaches were enslaved to the low statistical confidence because of lacking negative experimental verified profiles. Considering that, Chen and Yan [12] come up with a Regularized Least Squares (RLS) based method which is a semi-supervised classification approach, it can tackle the problem without utilizing negative samples.

Network-based methods generally sorted the prediction results, selected some of the top candidate entries, and then recommended them to biologists. These models assumed that those similar lncRNAs/miRNAs are prone to have common associations related to particular diseases[13–15]. Considering this acknowledgment, Sun et al. [16] proposed a computational method using globally optimized strategy named RWRlncD and Chen [17] utilized similarity construction on network to measure the relationship between miRNA and disease. Collected miRNA-disease (lncRNA-disease) connection and the directed acyclic graph (DAG) regarding to disease annotation from online web MeSH (https://www.nlm.nih.gov/), some researchers tried to construct disease-disease similarity network. Then these data was used to infer more positive disease-related miRNAs/lncRNAs[18], trying to boost the model performance. After obtaining all the similarity and association matrix, researchers implemented network-based algorithms such as Random Walk with Restart[16] or KATZ method[19] to get predicted disease-related miRNAs/lncRNAs[20]. There are other recent deep learning based methods like TLNPMD[21], MINIMIDA[22], etc. that utilized convolutional network for feature extraction and association inference, but they suffer from high computational cost due to large trainable parameters.

Due to the similar relationship among miRNAs, diseases and circRNAs, all the methods previously used by researchers could apply to circRNA-disease association prediction provided that similar interaction data is obtained. However, considering that there is not enough biological evidence directly showing the links between diseases and circRNAs, other information is of necessity to help infer the circRNA-disease association. It is worth noticing that Memczak et al. [2] has collected 1953 presumed human circRNAs along with their locations from genomic regions and predicted miRNA seed correspondence, which inspires us that it could probably been made good use of. Utilizing this information, Ghosal et al. [5] constructed a database Circ2Traits which provided miRNA-related interactions with circRNAs and diseases. Gladly, disease related miRNAs and miRNA related circRNAs data could be retrieved from Jiang et al. [7] and Memczak et al. [2]. So in this paper, we are searching for a proper way to infer circRNA-disease association through computational model and we managed to propose a deep ensemble framework to explore complex features from circRNA and disease information and then implemented a way to conjecture the paired association. In the view of scarcity of biological validated circRNA-disease association, We attempted to constructed three potential circRNA-disease association matrices as candidate labels and developed a ensemble model based on two auto-encoders with fully-connected network architecture for inferring the association on circRNAs and diseases. The prediction result was validated by several biological experiments already published in previous literature.

The overall paper layout is constructed as follows: (1) The METHODS illustrates the overall design related to this framework. (2) RESULTS section performs several benchmarking comparisons against other competitors and case studies. (3) DISCUSSION section conclude the whole result. The main contributions of this paper are listed here: (1) The construction of circRNAs and diseases similarity was based on ternary interactions, i.e. miRNA-disease-circRNA associations instead of pairwise analysis. (2) A deep learning model was built to generate hidden biological features using original similarity information instead of integrating multiple data sources and directly implementing features to different models. (3) Case studies was performed to further validated the prediction results on real circRNA-disease connection.

## II. Methods

### A. Human circRNA-miRNA associations

Circ2Traits database has collected 4583 verified human circRNA-miRNA associations including 101 miRNAs and 923 circRNAs and 765 paired associations consisting 101 miRNAs and 104 diseases. Using interaction data, we further constructed two adjacency matrix and **M**_*mc*_ and **M**_*md*_ to represent miRNA related associations with both circRNA and disease. For example, if miRNA *i* shown as *m*(*i*) is known to be related with disease *j* shown as *d*(*j*) from the database Circ2Traits, which was verified existed in HMDD[6], the score of *m_md_* (*i,j*) is 1 otherwise 0. Similarly, if miRNA *i* shown as *m*(*i*) is paired with circRNA *c*(*k*), the score of *m_mc_*(*i,k*) is set to 1 otherwise 0.

### B. Improved TAM method for circRNA-disease association matrix

Potential circRNA-disease associations were acquired using a statistical enrichment analysis method called TAM by performing hypergeometric test[23]. We calculated the statistical significance of the enrichment of each circRNAs interacting with disease related miRNAs. For each circRNA that is associated with miRNAs, if it is associated with a specific disease, the significance value denoting the likelihood of the circRNA to be interacted with one disease is calculated as follows:

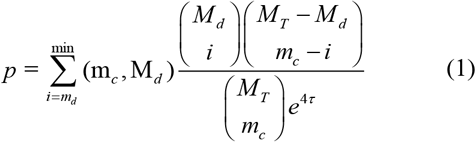

Where, *M_T_* stands for all the number of miRNAs, m_*c*_ refers to the number of miRNAs which have interaction with the specific circRNA, *M_d_* denotes the count number of miRNAs related to disease d, md represents the number of miRNAs related to disease d that associated to the specific circRNA and τ is a binary number denotes whether the circRNA is related to disease-related SNP. The SNP information was obtained by mapping the variation locations from human DNA/RNA sequence, which can be retrieved from dbSNP database, into the locations of circRNAs[5]. Those disease related SNPs that were mapped to the loci of circRNAs were considered as potential disease related circRNAs and its τ is 1 otherwise 0.

The P-values were finally adjusted using Bonferroni correction(using python function multipletest)[24] and those with p-value less than 0.05 is considered as likely disease associated circRNAs because it is highly possible significant p-value indicates that the correlation between circRNA and disease is most likely reliable. The circRNA-disease association matrix is represented as 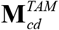.

### C. Network based matrix projection for circRNA-disease association matrix

Using two matrix and **M**_*mc*_ and **M_*md*_**, we could construct circRNA-disease association matrix, providing that if one miRNA is likely to be associated with a disease then any miRNA-related circRNA is likely related to this disease due to the functional mechanism of circRNA-miRNA interaction. The circRNA-disease matrix 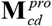 is defined as follows:

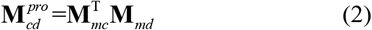

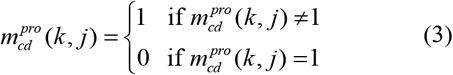

where 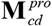 is a matrix whose entity 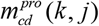 denotes the association score between circRNA *k* shown as *c*(*k*) and disease *d*(*j*). The value of 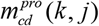 is 1 indicating circRNA *c*(*k*) is related to disease *d*(*j*) otherwise 0.

Furthermore, we constructed a mix label matrix 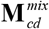 which used the intersection of two matrices 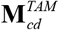 and 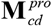, i.e. if item (*i,j*) is 1 in both matrices, 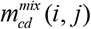 is 1 otherwise 0.

### D. Gaussian kernel based interaction similarity profile for circRNAs and diseases

Providing that functional similar miRNAs are prone to interact with another circRNAs with similar regulated function and vice versa. So we adopted a widely used technique by using Gaussian kernel to calculate circRNA similarity *KC* between circRNAs *c*(*i*) and *c*(*j*) and disease similarity *KC* between disease pair *d*(*i*) and *d*(*j*) from the known miRNA-circRNA association and miRNA-disease data respectively, which can be descibed in the equations below:

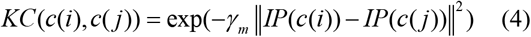

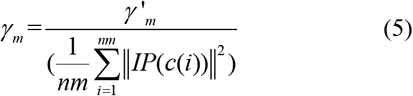

where we included *γ_m_* to control the bandwidth w.r.t. the kernel, this is calculated through introducing another new bandwidth parameter *γ’_m_*, normalizing it by calculating the average number of associations with miRNAs per circRNA. Here we set *χ’_m_* = 1 for simplicity.

### E. Disease similarity construction using Semantic feature

Individual disease can be described as an item in a Directed Acyclic Graph (DAG) and each item can be acquired through MeSH database, website is http://www.ncbi.nlm.n-ih.gov. *DAG*(*D*) = (*D, T*(*D*), *E*(*D*)) can be used to stand for the disease D, where *T*(*D*) stands for the node set that includes node D itself and its corresponding ancestor (or called upper) nodes, *E*(*D*) accounts for the edge set linking from parent to child nodes[25]. Every disease has only one or multiple MeSH ID that consecutively defines its location appeared in MeSH graph. We adopted the cosine similarity method used in Ding et al. [26], which was developed to improve the reliability of disease similarity calculation by using semantic feature. For each disease item, we got its feature vector v based on its hierarchical DAG information, and then computed the similarity semantic score of *d*(*i*) and *d*(*j*) by cosine function:

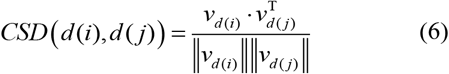

where *v*_*d*(*i*)_ and *v*_*d*(*j*)_ are the feature vector of disease *d*(*i*) and *d*(*j*), respectively.

Integrated disease similarity matrix *S_d_* was finally computed using either disease Gaussian kernel based similarity profile, or cosine similarity of disease semantic features respectively.

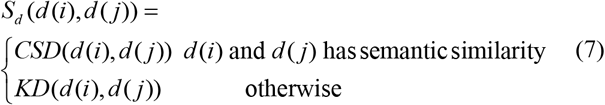

### F. DeepInteract model

In this study, we presented an ensemble deep learning framework, DeepInteract, to predict circRNA-disease association. DeepInteract, contains two main sub-modules. First, it extracted circRNA and disease similarity features separately by applying two stacked auto-encoders[27]. Secondly, two separated features were merged, combined with a P-value we got from Equation 1 (see Methods) to form a new feature vector and then fed into another auto-encoder with a softmax layer to cluster the circRNA into disease-related one or non-disease-related one. The overall flowchart is depicted in Fig.1. Python code and dataset are all available at https://github.com/sperfu/DeepInteract.

**Fig. 1.**
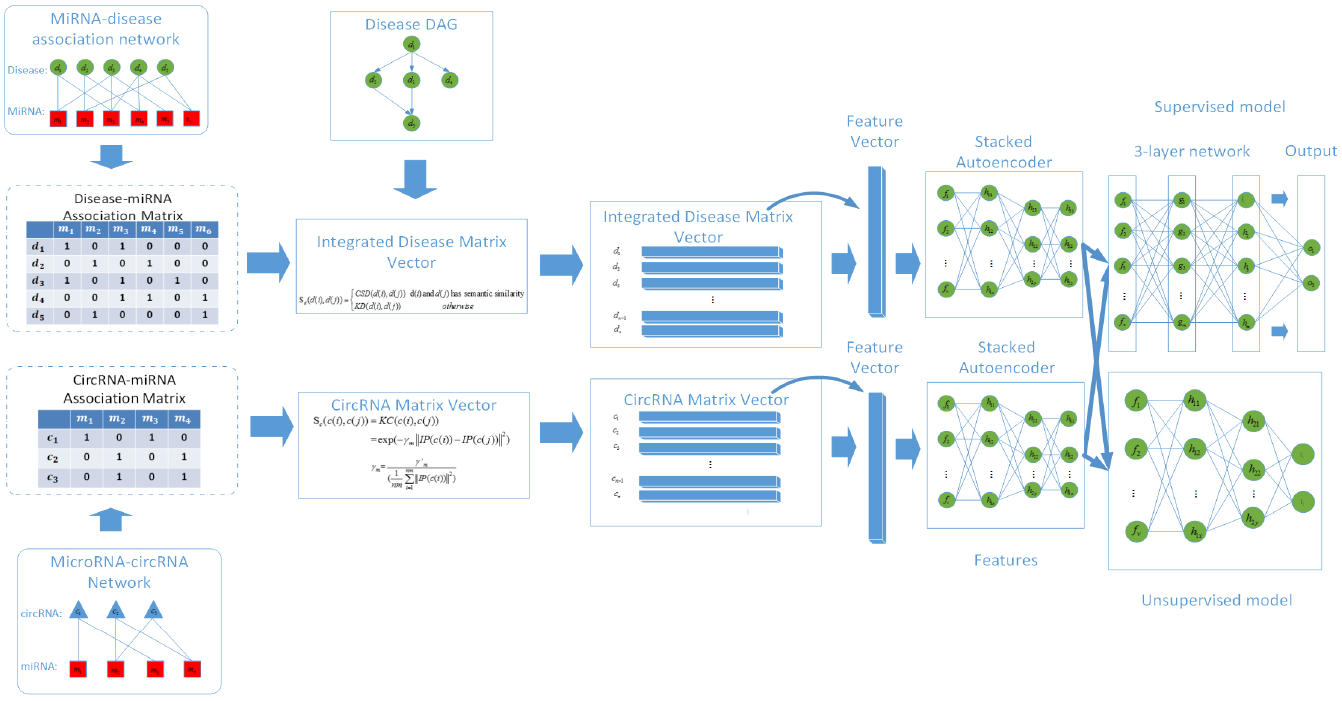
The flowchart of proposed DeepInteract. The two similarity matrices were separately computed using miRNA related associations with circRNA and disease. we also adopted DAG information for disease similarity calculation. We fed the two input vectors into two separated stacked autoencoders separately to learn hidden features and then concatenated them. In the model, it finally adopted a 3-layer fully-connected network to finally predict the associations between circRNAs and diseases.

To utilize the label matrix as we mentioned previously, we also developed a deep neuron network consisting of 3 layers of MLP network connecting to the output of two autoencoders to finally predict the association score between a circRNA and a disease. The overall parameter counts of DeepInteract are 581,873, which are relatively less than most CNN or RNN based models. The overall proposed framework of DeepInteract is depicted in Fig. 1.

### G. Stacked auto-encoder

After retrieving the similarity matrix of circRNA and disease, we spliced the circRNA similarity matrix into *n_c_* rows corresponding to *n_c_* different circRNA samples and cut disease similarity matrix into *n_d_* rows corresponding to *n_d_* various diseases, where every single line in the two matrices corresponds to a single sample of a circRNA or a specific disease. These samples were separately fed into the first part of deep network consisting of multiple layers. The *n_c_* circRNA samples were put into one auto-encoder while the *n_d_* disease samples were fed into another auto-encoder to learn features separately.

The input data is the similarity feature vector with d-dimension, it is mapped to y:

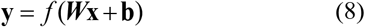

where *f* is the chosen activation function which projects the linear transformation w.r.t ***x*** to a non-linear feature. After that, the embedding ***y*** used the same procedure to compute ***z***, whose shape is the same as ***x***, showed as below:

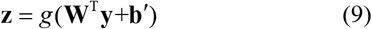

where *g* represents another function similar to *f*.

Here we used greedy search strategy, which use layer-wise learning by focusing on one layer while keeping other layers’ parameters freezing and chose mean squared error (MSE) between **x** and **z**, this error can be minimized by choosing a frequently used stochastic gradient descent(SGD) method. We chose dropout layers to help alleviate over-fitting problem through randomly drop out several neuron units with the rate setting to 0.5. After the first stage, we obtained the extracted high-order features passing to next part of training and final prediction.

### H. Deep Neuron Network

The two separate outputs generated by two auto-encoders were then merged to form into an independent sample feature vector. Every single line in the similarity matrix represented each specific circRNA or disease, and every single circRNA feature vector was concatenated to each disease feature vector separately, so there were *n_c_ ·n_d_* independent merged samples altogether, the concatenated sample was labeled by using corresponding pair-wise position status in the circRNA-disease interaction label matrix we previously got. Here, we used three association datasets, 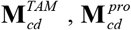 and 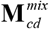 as label matrices separately to evaluate their performance. The final concatenated feature vector was then put into a feedforward network, which comprises a three-layer fully connected networks. And sigmoid function is chosen for final prediction. Cross-entropy cost function was finally chose to calculate the overall loss called C defined as follows.

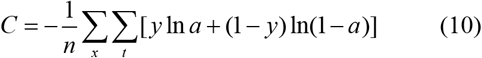

where *t* indicates the index of different labels, and *x* refers to each of our training sample, *y* is ground truth label and *a* is output prediction for 0 or 1 label given *x*.

ADADELTA optimized method with minibatch size of 200 is selected to calculate the training loss. NVIDIA Tesla graphic K80 GPU card was utilized to learn our model.

### I. Other composite models

For further comparing the performance of different models, then we designed and utilized a stacked auto-encoder(SAE) with Adaboost framework, we called it SAE-ADA, as another comparative method. We even used the latter part of DeepInteract, that is utilizing three layers Deep Neuron Network only to construct the classifier and using the similarity data as raw input (RAW-DNN). These two alternative deep model classifiers were compared with DeepInteract which used SAE as front part and implemented three layers deep network as the latter part (SAE-DNN). All these models performances were shown in the Result section.

## III. Results

### A. Inferring human circRNA-disease association via classifiers using five-fold and Leave-one-disease-out cross validation

In order to predict circRNA-disease association via classification, it is crucial to find a proper circRNA-disease association matrix as label matrix, so we constructed three association matrices according to previous research. First of all, we used TAM method[23], which was a statistical enrichment analysis method previously used to predict potential lncRNA-associated disease. We first adopted a revised TAM method to obtain the circRNA-disease association matrix 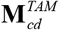 and then we utilized another matrix projection method to get an alternative association matrix 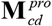 as comparison (see Method). Furthermore, we also used the intersection of two matrices denoting as a mixed matrix 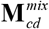 as association matrix. These three matrices were separately regarded as three independent label matrices corresponding to each circRNA-disease pairs. For comparison, we also applied another two deep learning models and multiple kinds of network-based algorithms to predict circRNA-disease association such as Regularized Least Squares based method[12, 28], Random Walk with Restart method[29], KATZ[19], Label Propagation[30] etc. We first used our model DeepInteract to evaluate on three label matrix, i.e., 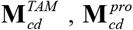 and 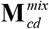. Five-fold cross validation (5-CV) was chosen to fairly evaluate the prediction models. For three label matrices 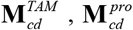 and 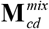, DeepInteract gained average AUC value in 5-CV of 0.9906, 0.9959, 0.9910 with the s.d.(i.e. standard deviation) of 0.002, 0.001, and 0.005, and the mean AUPR scores were 0.8630,0.9889,0.8778 with the s.d. of 0.023,0.001,0.016 respectively. Furthermore, we chose another measurement recall value to evaluate the model ability of retrieving the positive samples. As it was showed in Table I, DeepInteract has obtained 0.7521, 0.9205, and 0.7714 with the s.d. of 0.023, 0.009, and 0.039. When we compared our predicting result with some already validated experiments (see case studies), for example, gastric cancer, breast cancer etc., we found that using 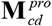 as label matrix could retrieve more biological confirmed circRNA-disease association. We then chose a more reliable one 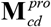 for later comparison.

**TABLE I.** Results on the three label matrices using DeepInteract.

| Label Matrix | AUC | AUPR | Recall |
| --- | --- | --- | --- |
| $\mathbf{M}_{cd}^{TAM}$ | 0.9906±0.002 | 0.8630±0.023 | 0.7521±0.023 |
| $\mathbf{M}_{cd}^{pro}$ | 0.9959*±0.001 | 0.9889*±0.001 | 0.9205*±0.009 |
| $\mathbf{M}_{cd}^{mix}$ | 0.9910±0.005 | 0.8778±0.016 | 0.7714±0.039 |

The three scores are calculated in the table. The \* mark refers the highest AUC/AUPR/Recall score. Overall, $\mathbf{M}_{cd}^{pro}$ has achieved best performance under three measurements.

To further explain the reason of using the model structure like DeepInteract, we then integrated another two deep learning models as comparison which are illustrated in Methods. The same evaluation standard was used. Their AUC scores were all above 0.85, AUPR scores were above 0.7, which were listed in Table II. What is more, our DeepInteract model still achieved the best performance in these three deep models, showing great power in predicting circRNA-disease association. Moreover, DNN is inevitable in serving as the final classifier when our model is compared with SAE-ADA model.

**TABLE II.** Results on the circRNA-disease Datasets.

| Method | AUC | AUPR |
| --- | --- | --- |
| RLS | 0.6171±0.003 | 0.0857±0.002 |
| LRLS | 0.6965±0.007 | 0.1309±0.003 |
| KATZ | 0.7840±0.003 | 0.1357±0.003 |
| RWR | 0.7585±0.004 | 0.1771±0.003 |
| Label Propagation | 0.7585±0.004 | 0.1402±0.001 |
| DeepInteract | 0.9959*±0.001 | 0.9889*±0.001 |
| SAE+ADA | 0.8976±0.001 | 0.7319±0.004 |
| RAW+DNN | 0.9659±0.002 | 0.9266±0.003 |

The overall metric scores are shown. We choose \* to represent the highest score. Generally, three designed models outperform other five models.

To further illustrate the effectiveness of circRNA-disease association prediction result, we next implemented a unique strategy called leave-one-disease-out cross-validation (LODOCV) in three deep learning models. In each run of the method, for a chosen disease *d*(*i*), all the circRNAs pairs w.r.t disease *d*(*i*) were all regarded as test samples. We tried to uncover their labels using other information. As the result shown in Fig. 2 and Table III, DeepInteract has achieved the best AUC score and ROC curve.

**Fig. 2.**
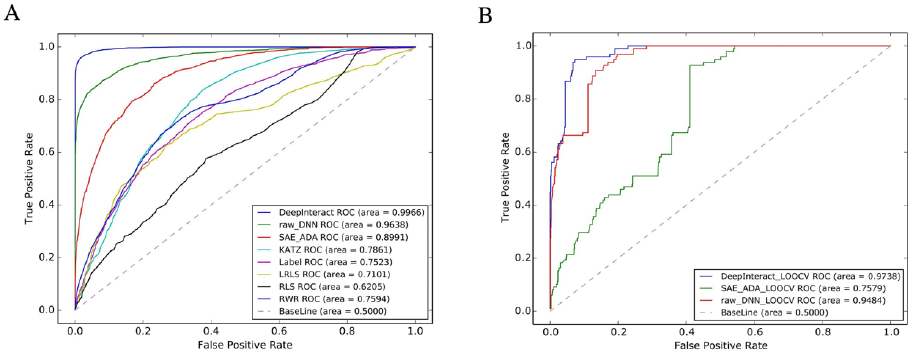
Prediction results on various compared methods. A) DeepInteract performed the best among eight different methods based on 5-CV test in ROC plot. B) DeepInteract achieves the best score in these three deep learning models in LODOCV test.

**TABLE III.** Results on the circRNA-disease datasets in LODOCV.

| Method | AUC | AUPR |
| --- | --- | --- |
| DeepInteract | 0.9036*±0.098 | 0.6984*±0.345 |
| SAE+ADA | 0.7752±0.165 | 0.4641±0.305 |
| RAW+DNN | 0.8296±0.179 | 0.6418±0.350 |

The overall metric scores are shown. We choose \* to represent the highest score. Generally, three designed models outperform other five models.

### B. Comparison with other methods

We also adopted other five models which were used in previous miRNA/lncRNA disease prediction models. 5-CV was implemented, Table II and Fig. 2 showed the prediction results of different models. From the result we could see that DeepInteract reached the best performance against all the compared methods with pairwise t-test p-value all less than 0.05. Generally, deep learning models demonstrated satisfactory and robust performance in circRNA-disease prediction using projection matrix as label matrix.

Overall, the performances of the designed deep learning based frameworks perform consistently better than the network based models, especially better in AUPR scores. Both the five-fold cross validation and LODOCV validation have achieved good performances. However, the result showed slightly lower scores in LODOCV than 5-CV because we left all circRNAs associated with one specific disease out in one turn. Predicting disease associated circRNAs without any known circRNA-disease information was much harder.

### C. Case studies

More biological evidence proved the connection between circRNA and disease through GO enrichment analysis[5]. Here, we presented several case studies about some vital diseases/cancers to prove the independent inferring ability of DeepInteract. The results are confirmed by searching for the biological literature for validation from Google Scholar.

Colorectal cancer(also known as bowel cancer), is one of the severely affected disease developed from the colon or rectum (parts of the large intestine). Specifically, hsa-circ-000737, hsa-circ-001569, and hsa-circ-001988 have reported aberrantly expressed in cancer of human-like colorectal carcinoma[4, 31, 32]. All these three circRNAs are all validated through our model. Especially, in the top 500 circRNAs our model predicted to be related with Colorectal cancer, 491 of which are confirmed in the benchmark dataset.

Similarly, other diseases such as gastric cancer, which is a disease developing from the lining of the stomach, may be related to circRNA hsa-circ-002059 and hsa-circ-000019, possibly indicating an unknown biomarker for gastric carcinoma[3, 33]. We were able to point out 188 circRNAs in the top 200 circRNA-gastric cancer associations using our model. And the prediction here may in some way provide some hints or clues for biological discovery so as to be helpful for the clinical and biomedical research.

## IV. Discussions

Substantial proofs have pointed out that circRNAs may consistently serve as an indicator or regulator of diseases development through regulating gene expression, quenching miRNAs activities and being the potential biomarkers, etc. Prediction of circRNA-disease associations could guide researchers to better understand the mechanism and inquire further reasons of different diseases. In our work, we presented a framework called DeepInteract to present circRNA-disease association prediction. First, circRNA-miRNA and miRNA-disease association matrix are both selected, we constructed three circRNA-disease association matrices as label matrices, circRNA functional similarity matrix and disease functional similarity matrix. Furthermore, we integrated disease similarities combining several designed similarity matrices. To begin with the model, the stacked autoencoder part of which was capable of capturing complex nonlinear features during learning phase using those similarity matrices as input and produced another representation of similarity regarding to every pair-wise sample, while the latter part of the model used another stacked auto-encoder or a 3-layer neuron network to predict association of circRNAs with diseases. Furthermore, both LODOCV and 5-CV were implemented, which had an average AUC of 0.9959, improving the second best model score up to almost 3 percent. Through the classifier model, we also showed some of the interactions that have even been confirmed by present papers such as has-circ-001569 related to colorectal cancer and hsa-circ-002059 associated with gastric cancer etc.

The performance of DeepInteract could attribute to some crucial factors. Firstly, various kinds of known datasets were adopted to explore the possible relations between different diseases and corresponding circRNAs. Secondly, DeepInteract was a deep ensemble learning method, which not only learned high level features itself through stacked autoencoders but also used multiple layer networks to construct powerful classifier or clustering tools and finally gained reliable results.

## V. Acknowledgment

We thank XiaMeng Wei for helpful discussion.

## VII. Author contributions

L.F. & Q.P. conceived the project, L.F. & H.D. designed the overall architecture, implemented the framework, and analyzed the result, L.F. & H.D. wrote the paper. H.D. and Y.W. polished the manuscript. Finally, all authors reviewed the final manuscript.

